# 2-amino-1,3-benzothiazole-6-carboxamide Preferentially Binds the Tandem Mismatch Motif r(UY:GA)

**DOI:** 10.1101/2020.06.11.146761

**Authors:** Andrew T. Chang, Lu Chen, Luo Song, Shuxing Zhang, Edward P. Nikonowicz

**Author notes:** Contribute equally (Andrew T. Chang, Lu Chen, Luo Song).

## Abstract

RNA helices are often punctuated with non-Watson-Crick features that can be the target of chemical compounds, but progress towards identifying small molecules specific for non-canonical elements has been slow. We have used a tandem UU:GA mismatch motif (5’-UG-3’:5’-AU-3’) embedded within the helix of an RNA hairpin as a model to identify compounds that bind the motif specifically. The three-dimensional structure of the RNA hairpin and its interaction with a small molecule compound identified through a virtual screen are presented. The G-A of the mismatch forms a sheared pair upon which the U-U base pair stacks. The hydrogen bond configuration of the U-U pair involves the O2 of the U adjacent to the G and the O4 of the U adjacent to the A. The G-A and U-U pairs are flanked by A-U and G-C base pairs, respectively, and the mismatch exhibits greater stability than when the motif is within the context of other flanking base pairs or when the 5’-3’ orientation of the G-A and U-U is swapped. Residual dipolar coupling constants were used to generate an ensemble of structures against which a virtual screen of 64,480 small molecules was performed to identify candidate compounds that the motif specifically binds. The tandem mismatch was found to be specific for one compound, 2-amino-1,3-benzothiazole-6-carboxamide, which binds with moderate affinity but extends the motif to include the flanking A-U and G-C base pairs. The finding that affinity for the UU:GA mismatch is flanking sequence dependent emphasizes the importance of motif context and potentially increases the number of small non-canonical features within RNA that can be specifically targeted by small molecules.

## Introduction

Non-coding RNA (ncRNA) molecules including microRNAs (aka, miRNAs) span a wide range of roles in biology including functioning as metabolic sensors, as scaffolds for assembly of large ribonucleoprotein complexes, and as direct suppressors of gene expression. To engage in their myriad functions, these molecules utilize both nucleotide sequence and shape (secondary and tertiary structure). Structured RNA molecules are primarily composed double helices that may contain conserved small structural motifs such as K-turns, adenine platforms, and loop E motifs. For RNAs with multiple helices, the arrangement of helices can be stabilized by tertiary interactions that include coaxial stacking, tetraloop:tetraloop-receptor interactions, and ribose zippers. RNA helices themselves are often interrupted by non-canonical elements such as non-Watson-Crick base pairs, bulged nucleotides, and internal loops. These elements, that can be central to an RNA molecule’s function, punctuate the regular A-form geometry of the helix with distinctive structural features or introduce local dynamic characteristics within an otherwise well-ordered helix.

Owing to their numerous cellular functions and their propensity to form secondary (and tertiary) structure, ncRNAs are attractive for their potential as targets of small therapeutic compounds ^*4, 5*^. Additionally, chemical compounds that are selective for specific structural features presented by an RNA molecule also can be versatile tools for RNA structure analysis and functional studies. The features recognized by many antibiotics that bind rRNA include regular helices and non-canonical elements and the same types of features are present in all RNAs. Further, multiple occurrences of bulged nucleotides and internal loops within a helix can establish a structural and dynamics pattern unique to the RNA molecule containing those features. Although non-canonical elements within regular helices offer clear opportunities for targeting specific ncRNAs, progress towards the identification of small molecules that selectively bind such non-standard features has only recently begun to gain momentum ^*6–10*^.

One lesser common irregular secondary structure element is the tandem UU:GA base pair mismatch. This mismatch motif occurs in the helices of large rRNA subunits of archaea, in many eukaryotic rRNA subunits in the region homologous to the bacterial ribosomal protein L11 binding site, within an HIV-1 intronic splicing silencer (ISS) stem-loop ^*11*^, and within the intermolecular helix formed between the anticodon arm of human tRNA^Lys3^ and the HIV A-rich Loop I adjacent to the HIV primer binding site ^*12*^. The UU:GA motif also forms part of the binding site of an aptamer of ribosomal protein S8 ^*13*^ and has been observed in the stems of at least 24 immature human miRNA molecules including miR-382 ^*14*^, miR-496 ^*15*^, and miR-508 ^*16*^. The tandem mismatch within the aptamer facilitates formation of a tertiary structure necessary for protein binding and may play a similar role in the rRNA L11 protein binding site ^*17*^. Structural studies of RNAs containing the UU:GA mismatch also suggest that the identity of the flanking base pairs may influence the structure and dynamics of the mismatch ^*11–13, 17*^.

We have used heteronuclear NMR spectroscopy to study the solution structure of a tandem UU:GA mismatch motif within an RNA helix and have identified a small molecule, ZN423 (IUPAC name: 2-amino-1,3-benzothiazole-6-carboxamide), that binds this motif. The mismatched nucleotides form U-U and sheared G-A base pairs with U_7_ O2 and U_22_ O4 serving as the hydrogen bond acceptors within the U-U base pair. ZN423 broadens the base resonances that flank the G-A pair that reflect chemical exchange on the intermediate timescale. Although the binding affinity of ZN423 to the RNA is moderate (K_D_ >100 uM), ZN423 exhibits selectivity for the motif. In addition, binding of ZN423 was found to be dependent on the identity of the base pairs flanking the UU:GA motif. Finally, although substitution of U7 with cytidine supports ZN423 binding, the U_7_-C_22_ variant does not bind ZN423. A structural model of the ZN423-RNA complex is presented.

## Materials and methods

All enzymes were purchased (Sigma) with the exception of T7 RNA polymerase which was prepared as described ^*18*^. Deoxyribonuclease I Type II, pyruvate kinase, adenylate kinase, and nucleotide monophosphate kinase were obtained as powders and dissolved into 15% glycerol, 1 mM dithiothreitol and 10 mM Tris-HCl, pH 7.4 and stored at −20 °C. Guanylate kinase and nuclease P1 were obtained as solutions and stored at −20 °C. Unlabeled 5’ nucleoside triphosphates (Sigma), phosphoenolpyruvate (potassium salt) (Bachem), and 99% [^15^N] ammonium sulfate and 99% [^13^C_6_]-glucose (Cambridge Isotope Labs) were obtained as powders. The identified hit compounds were purchased from ChemBridge Corporation.

### Preparation of RNA Samples

RNA I, (Figure 1), was prepared by in vitro transcription using T7 RNA polymerase and a synthetic DNA template ^*19*^. Unlabeled RNA molecules were prepared from 10 ml transcription reactions using 4 mM 5’-NTPs. Isotopically enriched RNA molecules were prepared from 6 ml transcription reactions using 3 mM ^13^C /^15^N-enriched 5’-NTPs as described ^*20*^. The RNA molecules were purified (20% w/v preparative PAGE), electroeluted (Schleicher & Schuell), and ethanol precipitated. The RNA was dissolved in 1.0 M NaCl, 20 mM potassium phosphate, pH 6.8, and 2.0 mM EDTA and dialyzed extensively against 10 mM NaCl, 5 mM potassium phosphate, pH 6.8, and 0.05 mM EDTA using a Centricon-3 concentrator (Amicon Inc.). The samples were diluted with buffer to a volume of 0.3 ml and lyophilized to powders. For experiments involving the non-exchangeable protons, the samples were exchanged twice from 99.9% D_2_O and dissolved in 0.33 ml of 99.96% D_2_O. For experiments involving the exchangeable protons, the samples were dissolved in 0.33 ml of 90% H_2_O/10% D_2_O. The samples contained 80-120 A_260_ O.D. units in 0.33 ml (0.9-1.4 mM). Isotope labeled sequence variants of RNA I (Figure S5) were prepared from 1.5 ml transcriptions.

**Figure 1.**
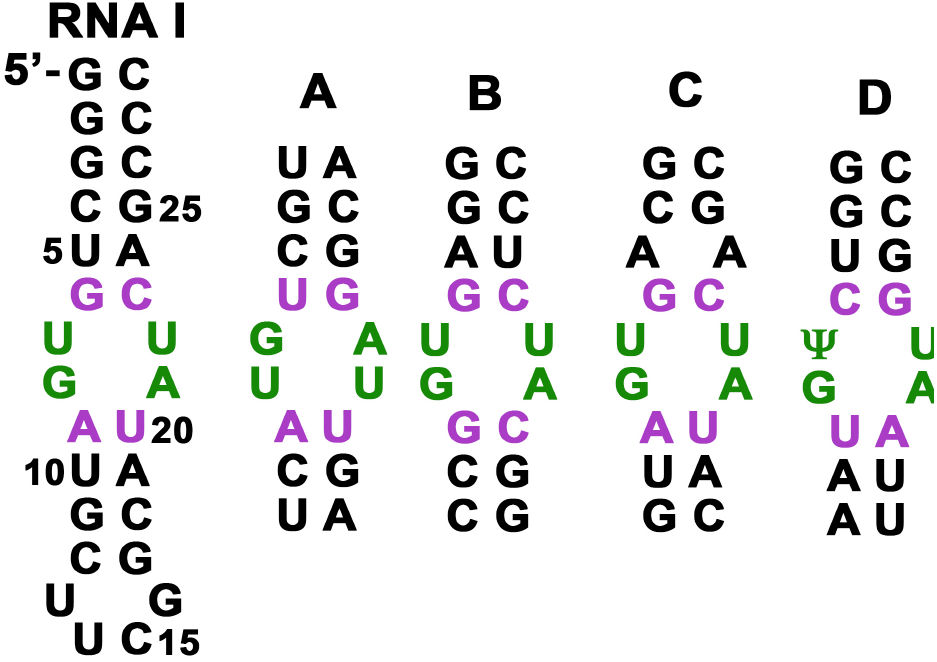
Secondary structures of RNA I and other tandem UU:GA mismatch-containing molecules. The three-dimensional structures of A-D have been solved previously. ^11,12,13,17^

### NMR Spectroscopy

Spectra were acquired on Varian Inova 500 MHz (^1^H-[^13^C, ^15^N, ^31^P] probe) and 600 MHz (^1^H-[^13^C, ^15^N] cryoprobe) spectrometers and were processed and analyzed using Felix 2007 (Felix NMR Inc., San Diego, CA). Two-dimensional (2D) ^13^C-^1^H HSQC spectra were collected to identify ^13^C-^1^H chemical shift correlations and to screen for RNA-compound interaction. Sugar spin systems were assigned using 3D HCCH-TOCSY (8 ms and 24 ms DIPSI-3 spin lock) experiments collected in D2O. Intra-residue base-sugar correlations were identified from H6-N1, H8-N9, and H1’-N1/N9 cross peaks in 2D H(C)N spectra. All pyrimidine correlations and several purine correlations were identified. Pyrimidine C2 and C4 resonances were assigned from H6-C2 and H5-C4 correlations using 2D H(CN)C and 2D CCH-COSY experiments ^*21, 22*^. 2D ^15^N-^1^H HSQC spectra optimized for 2-bond HN couplings were collected to identify purine N7 and adenine N1 and N3 resonances. Sequential assignments and distance constraints for the non-exchangeable resonances were obtained at 26 °C from 2D ^1^H - 1H NOESY spectra (t_m_ = 120, 160, and 320 ms) and 3D ^13^C-edited NOESY spectra (t_m_ = 160 and 320 ms). Assignments and distance constraints for the exchangeable resonances were obtained at 15 °C from 2D NOESY spectra (t_m_ = 160 and 320 ms) acquired in 90% ^1^H_2_O.

### Distance and torsion angle constraints

Interproton distances were estimated from cross peak intensities in 2D NOESY and 3D ^13^C-edited NOESY spectra. The covalently fixed pyrimidine H5-H6 distance (≈2.4 Å) and the conformationally restricted sugar H1’-H2’ distance (2.8-3.0 Å) give rise to the most intense peaks in the NOESY spectra and were used as a reference to assign upper bound distances to proton pairs based on the relative intensities of the corresponding cross peaks (qualitatively classified as strong, medium, weak, or very weak in the 120 ms and 160 ms mixing time spectra). None of the cross peaks are as intense as the pyrmidine H5-H6 peaks, thus upper bound distance constraints were set to 3.2, 4.2, 5.2, or 6.2 Å. Cross peaks observed only at mixing time 320 ms were classified as extremely weak and given 7.2 Å upper bound distance constraints to account for the possibility of spin diffusion. All distance constraints were given lower bounds of 1.8 Å. Only the intra-residue sugar-to-sugar constraints involving H5’ and H5’’ resonances included in the calculations are considered conformationally restrictive. Distance constraints involving exchangeable protons were estimated from 320 ms mixing time NOESY spectra and were classified as medium, weak, very weak, or extremely weak.

Watson-Crick base pairs were identified by observation of a significantly downfield shifted NH or NH2 proton resonance and the observation of strong G-C NH– NH2 or A-U H2–NH NOEs and by the chemical shifts of non-protonated base ^15^N and ^13^C carbonyl resonances. Hydrogen bonds were introduced as distance restraints of 2.9 ± 0.3 Å between donor and acceptor heavy atoms and 2.0 ± 0.2 Å between acceptor and hydrogen atoms. Ribose ring pucker and backbone dihedral constraints were derived from ^3^J_HH_, ^3^J_HP_, and ^3^J_CP_ couplings ^*23*^. Residues with ^3^J_H1’-H2’_ < 5 Hz and C3’ resonances between 70-74 ppm were constrained to C3’-*endo*. Ribose rings with ^3^J_H1’-H2’_≈5 Hz and with C3’ and C4’ resonances between 74-76 and 84-86 ppm, respectively, were left unconstrained. The angle *δ* was constrained as 85° ± 30° and 160° ± 30° for C3’-*endo* and C2’-*endo* sugars respectively. For residues 1 to 6, 9-12, 17-20, and 23-28, *γ* was constrained to the *gauche*^+^ conformation (60 ± 20°) ^*23*^. *γ* was left unconstrained for the internal loop residues. Dihedral angle restraints for the *β* and ε torsion angles were estimated from ^3^J_P-C2’/P-C4’_ couplings measured in 2D CECT-HCP ^*24*^ spectra. For stem residues, *β* was constrained to the *trans* conformation (180 ± 20°) if 3J_P-C4’_ was > 5 Hz. ε was constrained to the *trans* conformation (−150 ± 20°) for residues with ^3^J_P-C2’_ < 5 Hz and ^3^J_P-C4’_ > 5 Hz. *α* and *ζ* were constrained to −65 ± 20° for the stem residues 1 to 5, 10-12, 17-19, and 24-28. A down-field shifted ^31^P resonance is associated with the *trans* conformation of *α* or *ζ* and because no such shift is observed for any of the ^31^P resonances, *α* or *ζ* were loosely constrained to (0 ± 120°) which excludes the *trans* conformation for all other residues. Although all base 6/8-1’ intra-residue NOE cross peak intensities support the *anti* conformation about the glycosidic bond, no dihedral angle constraints were used for the angle *χ*.

^1^H-^13^C residual dipolar coupling constants (RDCs) were determined from the measured frequency difference between corresponding proton doublets in HSQC spectra acquired for isotropic and Pf1 phage-aligned samples. RDC values from base CH and ribose 1’ CH vectors were obtained in this way. The axial and rhombic terms were determined within XPLOR-NIH using an extensive grid search ^*25*^, and yielded values of D_*aH*_=−18.677 and R_*hH*_=0.153.

### NMR Structure calculations and refinement

An initial set of structures was calculated using a shortened version of the simulated annealing protocol (described below). A list of all proton pairs in the calculated structures closer than 5.0 Å (representing expected NOEs) was compared to the list of constraints. The NOESY spectra were then re-examined for predicted NOEs absent from the constraint list. In some cases, this allowed the unambiguous assignment of previously unidentified NOEs, but, in other cases, the predicted NOEs were obscured due to spectral overlap.

Structure refinement was carried out with simulated annealing and restrained molecular dynamics (rMD) calculations using XPLOR-NIH v2.44 with the RNA ff1 force field and potentials implemented ^*25, 26*^. Starting RNA coordinates were generated as a single stranded oligoribonucleotide with C3’-*endo* ring puckers for all ribose moieties using 3DNA ^*27*^. The structure calculations were performed in two stages. Beginning with the energy minimized starting coordinates, 50 structures were generated using 20 ps of rMD or 10000 steps, whichever is less, at 1200K with hydrogen bond, NOE-derived distance and base-pairing restraints. The system then was cooled to 25 K over 0.2 ps of rMD or 10000 steps, whichever is less. During this stage, RDC constraints and repulsive van der Waals forces were introduced into the system and the SANI force constraint used for RDCs was gradually increased from 0.010 kcal mol^−1^ Hz^−2^ to 1.000 kcal mol^−1^ Hz^−2^. Other force constants used for the calculations were increased—from 2 kcal mol^−1^ Å^−2^ to 30 kcal mol^−1^ Å^−2^ for the NOE and from 10 kcal mol^−1^ rad^−2^ to 100 kcal mol^−1^ rad^−2^ for the dihedral angle constraints. Each structure was then minimized with constraints at the end of the rMD. Eight structures were selected for the final refinement. The criteria for final structure selection included lowest energies, fewest constraint violations, and fewest predicted unobserved NOEs (^1^H pairs less than 3.5 Å apart, but no corresponding cross peak in the NOE spectra). A second round of rMD was performed on these structures using protocols similar to those used in the first round of structure calculation. The major difference was the starting temperature of 300 K followed by cooling to 25 K over 28 ps of rMD. Eight refined structures for each model were analyzed using XPLOR-NIH and UCSF Chimera ^*28*^.

### RNA structure ensemble generation

Coordinates from the energy minimized average NMR structure were used as the starting coordinates for a molecular dynamics (MD) simulation with GROMACS 5.05. The RNA molecule was solvated in TIP3P water and neutralized with sodium ions. After energy minimization, temperature and pressure of the system were equilibrated for 100 ps before the production run which continued for 100 ns. Snapshots were saved every 2 ps for further analysis.

The structure ensemble was generated by comparing the population average calculated RDC values with measured RDC values ^*29, 30*^. This method minimizes the number of conformers that satisfy all time-averaged RDC data and identifies a set of structures that represents unique and dominant populations across the entire RNA structure landscape. The structure ensemble was obtained using an in-house protocol termed *EnsembleGen*. The code and the detailed instruction for execution are freely available for download at https://imdlab.mdanderson.org/ressd/ressd.php. Combined with Relax 4.0.3 (http://wiki.nmr-relax.com/Relax_4.0.3), *EnsembleGen* uses one or more sets of RDC data to guide selection of RNA conformers from the pool of MD snapshots.

### Virtual screen for motif-selective compounds

*In silico* high-throughput screening was performed as reported previously ^*31*^ to identify small molecules potentially capable of specifically binding the UU:GA motif. Briefly, the ChemBridge diverse set (~50,080 compounds) and MayBridge (~14,400 compounds) were screened using GOLD docking software with GOLD Fitness scoring function ^*32*^. The top 10 poses for each compound were saved for rescoring using the rDock solv scoring function ^*33*^ and the top 3 poses were selected. These top poses were re-scored again using IMDLScore scoring function ^*31*^, and the top 10% compounds were selected for RNA binding studies.

Longer MD simulations were performed to elucidate the possible binding mechanism of a hit-RNA complex. The topology and charges of the small molecule were prepared using Gaussian09 at B3LYP/6-311++G(d,p) level of theory on the Texas Advanced Computing Center (TACC) cluster. The ff99bsc0 force field was used for the RNA and general AMBER force field (GAFF) for the hit prepared by ACPYPE ^*34*^. The complex was solvated in TIP3P water and neutralized with sodium ions. The simulation boxes were prepared so that no RNA or small molecule atom was closer than 14Å to the edge. The system was minimized and equilibrated for 2 ns before the production run which was conducted for 660 ns. Snapshots were saved every 20 ps for further analysis.

## Results and Discussion

### Spectral assignments

The non-exchangeable ^1^H and ^13^C resonances of RNA I were assigned using standard heteronuclear methods ^*35, 36*^. Most of the base and ribose resonances are resolved at 298 K and pH 6.8. None of the base or 1’ resonances were observed to be exchange broadened which would be indicative of intermediate time scale motions. Sequential assignment of non-exchangeable resonances was accomplished using 2D NOESY and 3D ^13^C-edited NOESY spectra. The sequential NOE connectivities are continuous in the base-1’ region (*τ*_mix_ = 160 ms) except at the U_7_-G_8_ step (Figure 2). The sequence specific resonance assignments are listed in Table S1.

**Figure 2.**
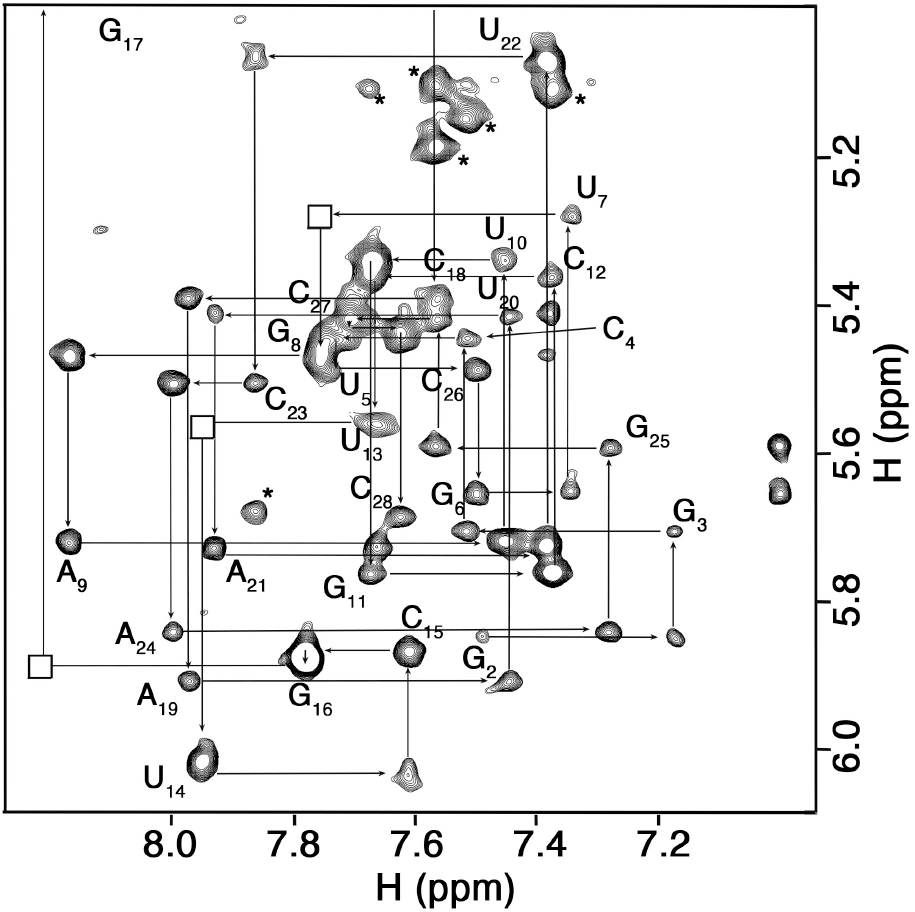
Base-1’ region of a pyrimidine C5 filtered 2D ^13^C HSQC-NOESY spectrum (t_m_ = 160 ms) showing the sequential walk. Residual pyrimidine H5-H6 cross peaks are indicated by asterisks. The U_7_-G_8_ and G_16_-G_17_ inter-residue base-H1’ cross peaks are weak, but present, in the corresponding 2D NOESY spectrum

The NH and cytidine NH_2_ resonances were assigned using ^1^H-^1^H NOESY and HNCCH experiments. The NH resonances of the stem base pairs, including the U-U base pair but not the G-A pair, yield NOE connectivities between neighboring base pairs. The G_8_ NH resonance was assigned based on ^1^H and ^15^N chemical shifts. The inter-nucleotide phosphate ^31^P resonances are clustered between −2.54 and −4.60 ppm, but were not sequence specifically assigned.

### NMR structure analysis

The solution structure of RNA I was calculated using a restrained molecular dynamics routine starting from 50 sets of randomized coordinates. The simulated annealing and refinement calculations were performed with 473 (305 NOE-derived and 68 hydrogen bond) conformationally restrictive distance constraints, 297 torsional angle constraints, and 51 RDC constraints (Table 1) leading to eight converged structures. The root mean square deviations (RMSDs) of the heavy atoms between the individual structures and the minimized mean structure is 0.194 Å.

**Table 1.** Summary of experimental distance and dihedral constraints and refinement statistics for RNA I.

| Constraint | RNA I |
| --- | --- |
| **NOE distance constraints** | **305** |
| Intra-residue<sup>a</sup> | 144 |
| Inter-residue | 161 |
| Mean number per residue | 10.9 |
| **NOE constraints by category** |  |
| Very strong (1.8 – 2.5 Å) | 4 |
| Strong (1.8 – 3.5 Å) | 11 |
| Medium (1.8 – 4.5 Å) | 104 |
| Weak (1.8 – 5.5 Å) | 105 |
| Very weak (1.8 – 6.5 Å) | 71 |
| **H-bond distance constraints** | **33** |
| **Dihedral angle constraints** | **297** |
| Ribose ring<sup>b</sup> | 72 |
| Backbone | 157 |
| Base pairs | 68 |
| Mean number per residue | 10.6 |
| **RDC constraints** | **51** |
| **Violations** | **0** |
| Average distance constraints > 0.3 Å<sup>c</sup> | 0 |
| Average dihedral constraints > 0.5°<sup>d</sup> | 0 |
| Average RDC constraints > 1.5 Hz | 0 |
| **RMSD from average structure<sup>e</sup>** |  |
| Heavy Atoms (Å) | 0.195 |

<sup>a</sup>Only conformationally restrictive constraints are included.
<sup>b</sup>Three torsion angles within each ribose ring were used to constrain the ring to either C2'-*endo* or C3'-*endo* conformation.
<sup>c</sup>A distance violation of 0.3 Å corresponds to 5.0 kcal energy penalty.
<sup>d</sup>A dihedral angle violation of 0.5° corresponds to 0.05 kcal energy penalty.
<sup>e</sup>Calculated against the minimized average structure.

Structure statistics are summarized in Table 1 and a representative structure is shown in Figure 3.

**Figure 3.**
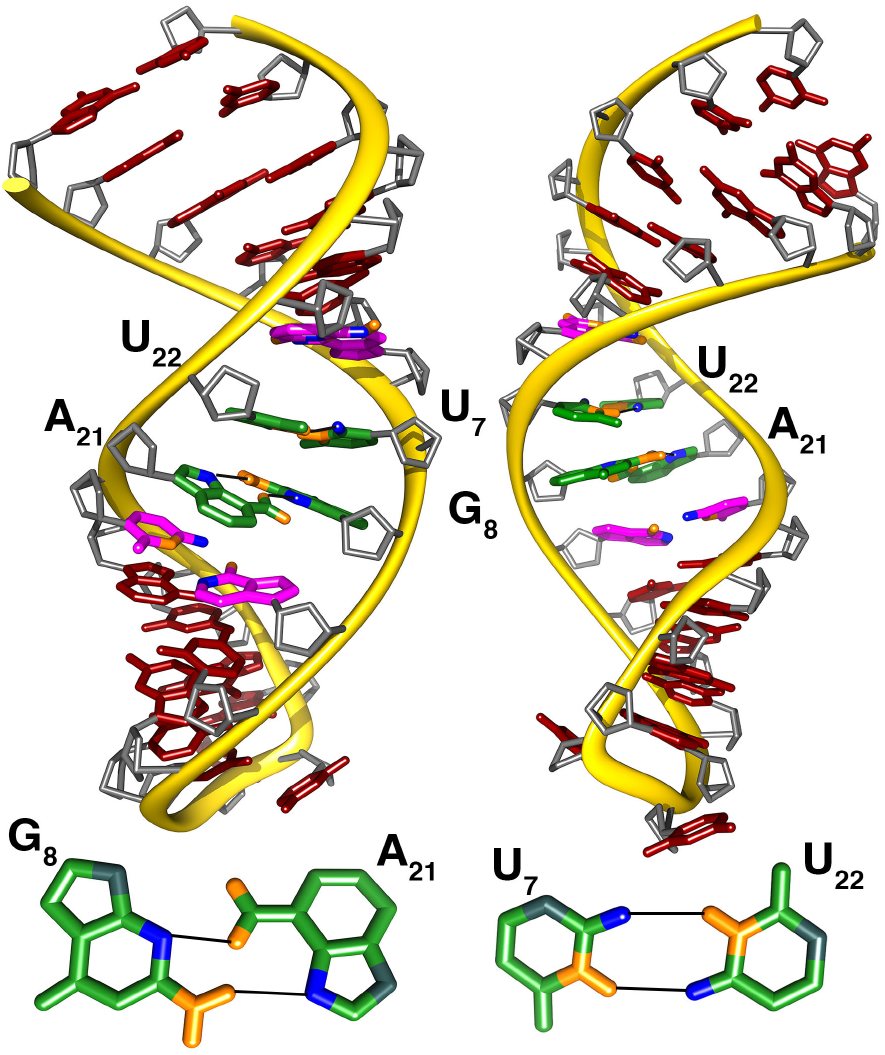
Minimized average solution NMR structure of RNA I with views into the major (left) and minor (right) grooves. The G-A and U-U mismatch bases shown below. The flanking A-U and G-C pairs are colored pink. The hydrogen bond acceptors (blue) within the mismatch are U_7_ O2, U_22_ O4, G_9_ N3, and A_21_ N7 and the amino and imino donor groups are colored orange.

### Structure around the UU:GA mismatch

The primary interactions of interest within the mismatch region (G_6_–A_9_ and A_19_– C_23_) are those that define the base pairing pattern of nucleotides. G_6_ and C_23_ and A_9_ and U_20_ form Watson-Crick pairs, although the U_20_ NH resonance displays solvent exchange broadening. The U_20_ N3 chemical shift of 156 ppm (identified through the U_20_ N3-H5 correlation) is consistent with an A-U base pair, but rapid exchange of the U_20_ NH resonance prevents confirmation of A_9_ as the base pair partner. Notably, the A_9_ C2 and H2 and U_20_ base 5 and 6 resonances do not display exchange broadening, pointing to increased solvent accessibility of the U_20_ imino proton rather than increased dynamics of the A_9_ or U_20_ bases as the cause undetected U_20_ NH resonance. Consistent with the H(N)CO spectrum (Figure S1), U_7_ and U_22_ form an asymmetric base pair where the hydrogen bond acceptors are O2 and O4 carbonyl oxygens on U_7_ and U_22_, respectively. The upfield shift of the U_22_ H1’ is consistent with the side by side (sheared) orientation of the G_8_ and A_21_ bases (Figure 3). Two NOEs of particular interest are the atypical A_21_ H2–G_8_ H1’ cross strand NOE and the unusually intense sequential A_21_ H2–U_22_ H1’ NOE cross peak (Figure 2). These interactions are consistent with the base of A_21_ stacked into the helix and is slid towards the minor groove away from the helix axis.

The sugar-phosphate backbone through the mismatch region of the stem has A-form geometry (Figure 3). The ribose puckers of G_6_–A_9_ and A_19_–C_23_ are uniformly C3’-*endo* (as they are throughout the stem). The phosphate backbone torsional angles among G_6_–A_9_ and A_19_–C_23_ are largely within regular A-form geometry except angles *α* and *β* between A_21_ and U_22_. *α* varies between *gauche*^+^ and *gauche*^−^ (rather than canonical *gauche*^−^) and *β* also tends to adopt either the *gauche*^+^ or *gauche*^−^conformation rather than the standard *trans* conformation. The ^31^P resonances are centered within the main cluster around −4.0 ppm, which is consistent with none of the *ζ*-torsional angles adopting the *trans* conformation.

The structure and dynamical properties of the of the tandem mismatch of U-U and G-A pairs have been examined in other contexts. In the context of G-C flanking sequences (Figure 1), the UU:GA internal loop appears ordered with intra-stem stacking of the uridine bases, inter-stem stacking of the adenine base and bulging of the G, but lacks regular base pairing. The arrangement of the internal loop is consistent with the NH resonances of the U bases that are significantly weakened by either solvent or chemical exchange ^*17*^. In a case where the order of the tandem mismatch is reversed and flanked by U-G and A-U base pairs (Figure 1), the G and A adopt the sheared conformation. However, the uridine bases are not hydrogen bonded generally oriented with their imino groups facing each other ^*11*^. The uridine NH resonances are not present in the spectrum, consistent with the absence of hydrogen bonding. A third structure of the UU:GA mismatch involves the substitution of pseudouridine (Ψ) for the 5’-U (5’-Ψ G). In this structure, with the mismatch flanked by C-G and U-A pairs (Figure 1), the Ψ-U also does not form hydrogen bonds. However, the G-A adopt the less common imino-imino conformation ^*12*^. Taken together, the structure of UU:GA motif is not only determined by the tandem pair of mismatched bases, but also by the identities of the flanking base pairs.

### Selection of the three-dimensional structure ensemble of RNA I

RNA molecules exhibit a wide range of motional time scales and amplitudes. Computational methods to probe the conformational states of RNA molecules have been used to identify structures capable of binding small molecules ^*29, 37*^. Thus, to explore the dynamics of RNA I and to generate ensemble structures that RNA I might sample, unconstrained molecular dynamics simulations with explicit solvent and counterions were performed using the minimized average coordinates of the solution structure. No significant conformational changes were observed through 100 ns of simulation. The RMSD between the starting structure and 50,000 snapshots from the simulation trajectory is illustrated in Figure S2.

Although the RMSDs of the RNA I coordinates during the simulation do not reveal dramatic structure changes, examination of individual snapshots shows that localized conformational fluctuations occur throughout RNA I, particularly around the mismatched bases. To address the structural flexibility of RNA I and improve the potential for success of the small molecule virtual screen, a structure ensemble was created by incorporating RDCs into the structure selection process using our in-house program *EnsembleGen*. Figure S3 compares the calculated and measured RDC values, with overall RMSD = 4.12 Hz. An ensemble of structures that reflects the unique and dominant conformations of nucleotides across the entire RNA structure landscape was constructed by determining the minimum number of RNA conformers that satisfy all time-averaged RDC data ^*38*^. The all heavy atom RMSDs between the minimized average NMR structure and the individual structures contained in the ensemble range 1.65-5.03 Å (Figure S2). The representative structures of RNA I from the dynamic ensemble are shown in Figure S2.

### Identification of small molecules specific for the UU:GA motif in RNA I

Previously, we examined computational docking tools to assess the predictive accuracy of experimental binding energies and established a protocol for RNA-small molecule docking ^*31*^. For each of the 20 conformers in the RNA I structure ensemble, virtual screening was performed against 64,480 commercially available small molecules from the ChemBridge and MayBridge databases. Using GOLD and rDock in a sequential docking procedure, a total of 16 compounds (Table S2) were selected for experimental validation based on their docking scores (top 100), size (MW < 400), and chemical similarity (Tanimoto Coefficient < 0.6). We also inspected the predicted binding poses for these compounds to ensure that the interactions between compounds and RNA are reasonable. Since no inhibitor has been reported as a reference, the criteria of reasonability is arbitrary, but we are especially interested in several features: docked sites (major groove vs. minor groove vs. intercalation between base pairs), interactions with based pairs vs. backbone, hydrogen bonding interactions, and aromatic stacking interactions.

NMR spectroscopy was used to evaluate the interaction of the 16 selected compounds with RNA I. Binding of a compound is expected to produce changes in the NH proton spectrum of RNA I. Although these compounds required the presence of 8-10% DMSO for solubility, concentrations of DMSO up to 15% caused no perturbations to the NH and base ^13^C-^1^H spectra of RNA I. This observation is consistent with previous NMR spectral studies of RNA utilizing DMSO as a co-solvent ^*39*^. Of the 16 compounds tested, two of them, 2-amino-1,3-benzothiazole-6-carboxamide (ZN423) and 5,7-dimethyl-1,4-dihydro-2,3-quinoxalinedione (ZN449), were found to perturb the NH spectrum, with the former causing substantial changes to the U_22_ and G_6_ peaks (Figure S4). ZN423 also leads to changes in the non-exchangeable ^13^C-^1^H HSQC spectrum (Figure 4). The base 2 and 8 resonances of A_9_ and base 6 resonance of U_7_ are exchange broadened beyond detection by ZN423 and the base 8 resonance of G_8_ is significantly weakened. Also, the G_8_ and U_20_ 1’ are exchange broadened by ZN423 (Fig 4).

**Figure 4.**
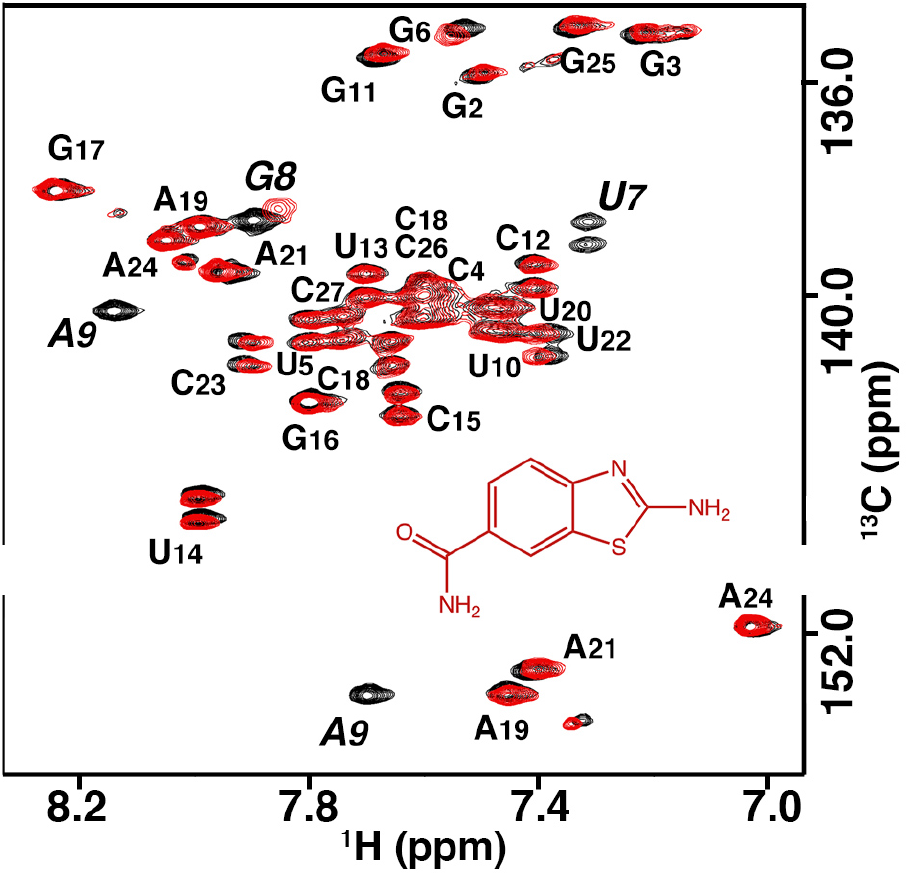
Base region of 13C-1H HSQC of RNA I in the absence (black) and presence (red) of ZN423. In the presence of ZN423, the U_7_ H6, A_9_ H8, and A_9_ H2 resonances are broadened beyond detection and the G_8_ H8 is broadened and substantially weakened. The chemical structure of ZN423 also is shown.

To examine the selectivity of ZN423 for the UU:GA motif, RNA spectra of hairpins containing substitutions of nucleotides within and flanking the mismatch loop were screened in the presence of ZN423. The base CH spectrum of a U_7_C variant that introduces a C_7_–U_22_ mismatch displays the same exchange broadening pattern as the native molecule except that the G_8_ base resonance is completely exchange broadened in the U_7_C variant. A U_7_A variant displays no spectral changes in the presence of ZN423, but interestingly destabilizes the A_9_-U_20_ and G_8_-A_21_ pairs, as evidenced by exchange broadening of the A_9_, U_20_, and A_21_ base resonances. Similarly, the spectra of U_22_C and A_21_C variants displayed no changes when ZN423 was added.

The effect of changing the flanking sequences was also examined. No evidence of binding could be observed after swapping the A_9_-U_20_ base pair with G_9_-C_20_ and C_9_-G_20_. Similarly, binding is significantly diminished when the U_10_-A_19_ pair is replaced with G_10_-C_19_. The U_5_ was replaced with an adenine nucleotide to create an A_5_-A_24_ mismatch and decrease the stem stability without completely disrupting the U_7_-U_22_ pair. This modification also appears to decrease the binding (diminished resonance broadening) of compound ZN423. The results of the various substitutions on ZN423 interaction with the RNA are summarized in Figure S5.

### Modeling of RNA-ligand structures

The binding affinity between ZN423 and RNA I is moderate (>0.1 mM), resulting in chemical exchange on the intermediate time scale and preventing solution NMR structure determination of the complex. No NOEs between RNA I and ZN423 could be identified in long or short mixing time NOESY spectra. However, the H5 of ZN423 displays exchange broadening in the presence of RNA I whereas H7 and H8 appear unchanged, suggesting a more proximal relationship of one edge of ZN423 with RNA I (Figure S6). Thus, to better understand the interaction of ZN423 with the UU:GA internal loop, unconstrained MD simulations of RNA I with ZN423 were performed. Figure 5A shows the smoothed trajectory of compound ZN423 over the ~660 ns simulation in Each stable state is assigned an ID, with its 3D structure illustrated in (B). explicit solvent. Interestingly, we observed a full binding cycle of ZN423’s association and dissociation with the UU:GA motif (Figure 5A-B) and identified six states in which ZN423 interacts with the RNA (Figure 5B). The compound quickly bound the minor groove of RNA I at the UU:GA motif, with a center of mass (COM) distance (Figure 5C) of 10Å between UU:GA and ZN423 by forming stable interactions (described below) for ~100 ns. The ZN423 molecule dissociated from RNA I and periodically (at 150 ns, 230 ns, and 300 ns) stacked against the terminal nucleotides. The compound also interacted with the major groove in a non-specific and transient manner (at ~450ns). Finally, it again bound to the minor groove of the UU:GA motif. The two binding modes in the minor groove are nearly identical, with the latter having a COM distance of 8Å. The residence time for the second minor groove binding event is ~80ns, followed by a second binding cycle (Figure 5A).

**Figure 5.**
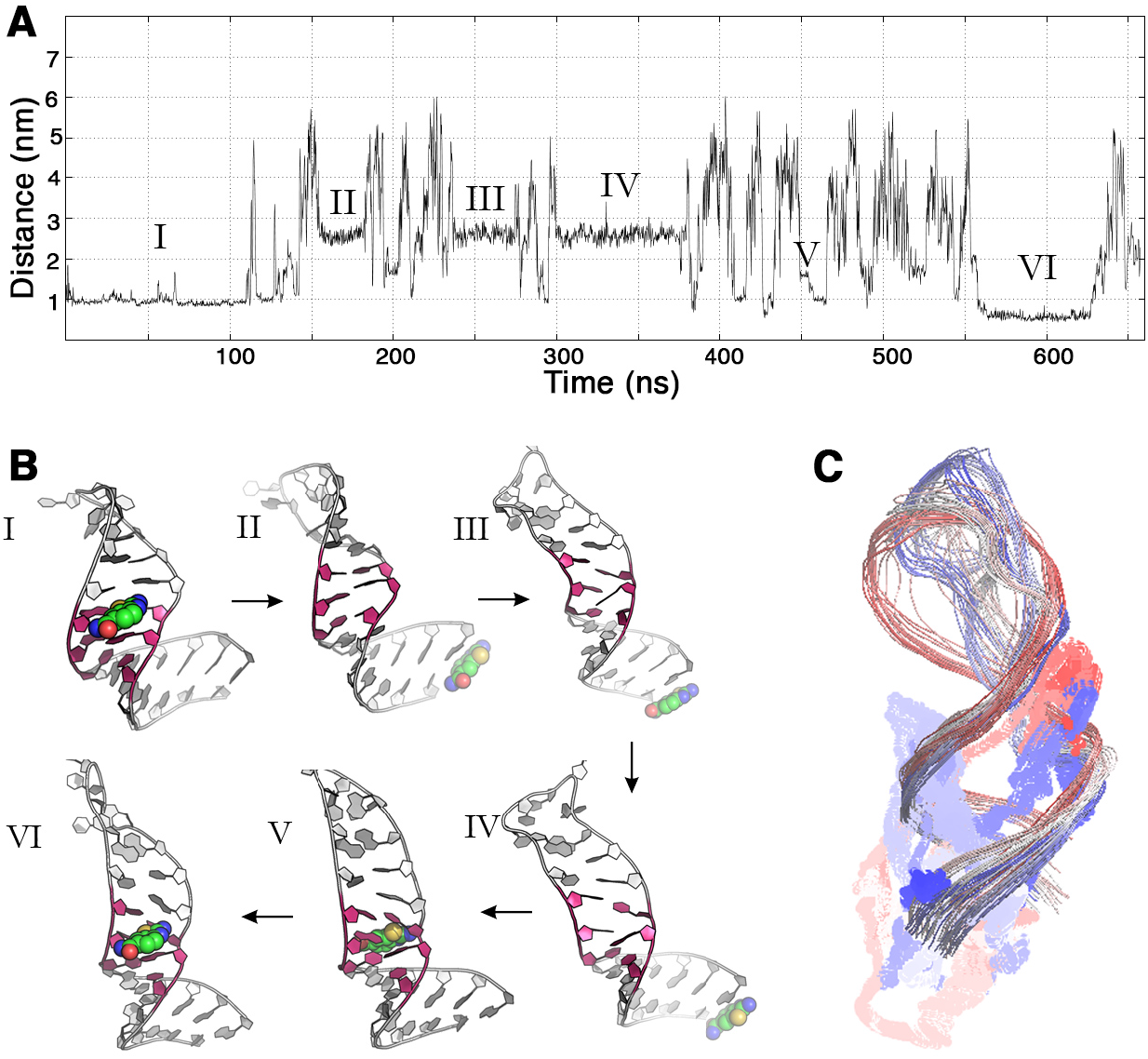
MD simulations of compound ZN423 binding to UU:GA motif. (A) Smoothed trajectory of RNA I and ZN423 over 660ns simulation. The color ranges from red to blue, denoting the time-dependent evolution of the complex structure from 0ns to 660ns. (B) Representative structures from the MD simulation. (C) Center of mass distance between the UU:GA motif and compound ZN423.

In both complexes involving the UU:GA motif, ZN423 binds to the minor groove defined by G_8_, A_9_, U_20_, and A_21_ (Figure 6A). The benzothiazole moiety stacks on A_21_, and the amine group interacts with U_20_ O2 to form an intermolecular H-bond (Figure 6A and Figure 6C). The sheared G-A base pair exhibits a propeller twist of 40° with the binding of ZN423 (Figure 6A). The moderate binding affinity (>0.1 mM) and the positioning of ZN423 in the minor groove causes exchange broadening of several resonances. The A_9_ H2 and G_8_ H1’ are proximal to the amine and benzene H5, respectively, of ZN423, which is consistent with the exchange broadening pattern. As ZN423 binds, the propeller twist of the G-A pair increases significantly, altering the local magnetic environments, and therefore chemical shifts, of several proximal protons including U_7_ H1’, U_22_ H6, G_8_ H8 and A_9_ H8.

**Figure 6.**
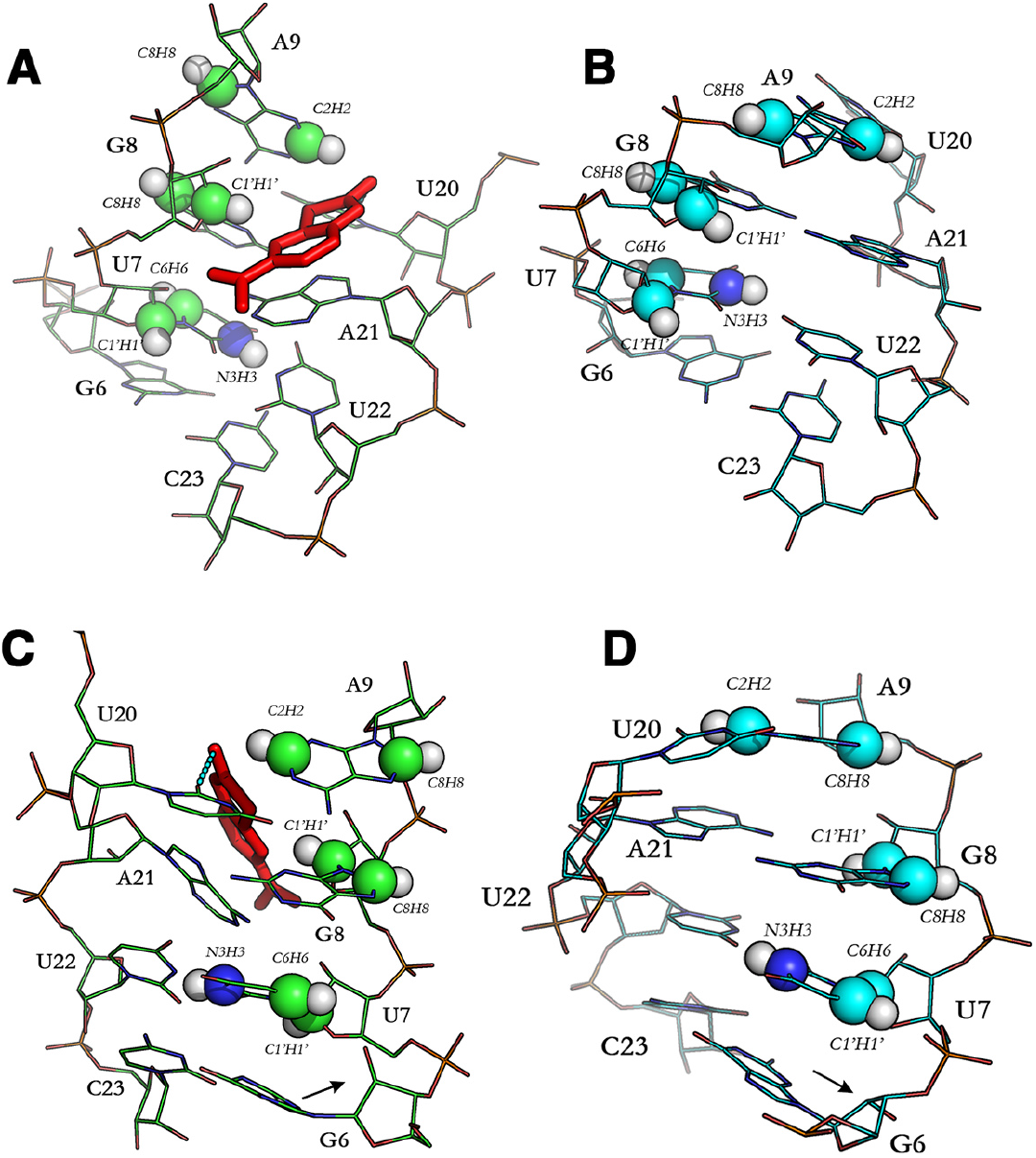
The 3D model of ZN423 binding to UU:GA motif. ZN423 is shown in red sticks, and the RNA atoms altered by the binding are shown in sphere. The changes of sugar puckering are highlighted with arrows. (A) Minor groove view of ZN423-bound complex structure. (B) Minor groove view of unbound RNA structure. (C) Major groove view of ZN423-bound complex structure. (D) Major groove view of unbound RNA structure.

Moreover, as illustrated by Figure 7, destabilization of the two base pairs next to the GA pair is an essential step before ZN423 binds to the UU:GA motif. In comparison, the base pair stability at the +3 position does not correlate with the compound binding event. Thus, the MD simulation suggests the importance of flexibility on the GA side of the mismatch for binding affinity and this is consistent with the flanking sequence dependence on ZN423 binding observed by NMR (Figure S5).

**Figure 7.**
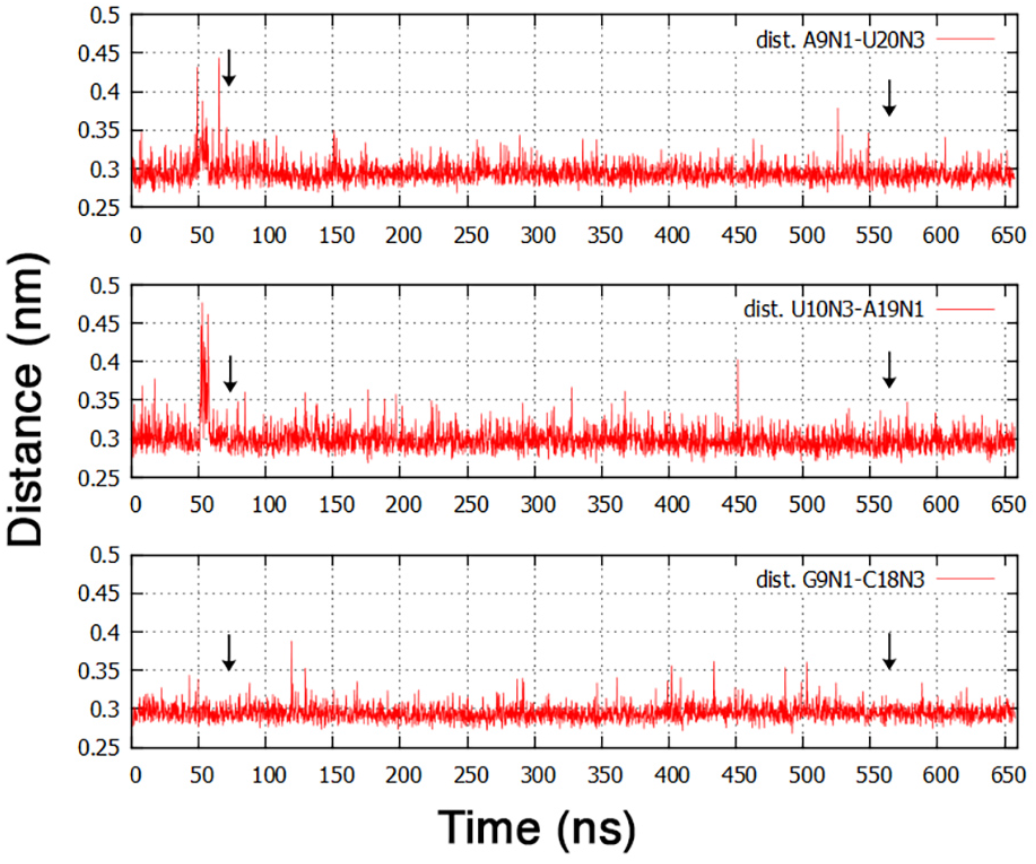
Base pair flexibility in the context of UU:GA motif. Time-dependent distance between A_9_ N1-U_20_ N3, U_10_ N3-A_19_ N1, and G_9_ N1-C_18_ N3, which denote the three base pairs from the GA base pair. The arrows highlighted the time point that ZN423 started to associate with UU:GA motif.

We further assessed the AN423-RNA binding model indicated by the MD simulations experimentally using four derivatives of ZN423 (Table 2). The compounds were subjected to *in silico* docking analysis and titration with RNA I monitored by NMR. Any substitution at the R1, R2 or R3 positions failed to cause changes in the NMR spectra at concentration as high as 0.2mM (Table 2). This apparent lack of interaction is consistent with our structure models in which the amine group (R1) forms a hydrogen bond with U_20_ and the carboxamide group (R3) forms polar contacts with the G_8_ ribose ring such that any hydrophobic substitution would be expected to impair or abolish the binding. Removal of the carboxamide from R3 and substitution at R2 with a hydrazide moiety was also found to be detrimental to binding. Although the structure model suggests that R2 is exposed to the solvent and thus is expected to minimally contribute to binding, the hydrazide group cannot substitute for the simultaneous loss of the R3 carboxamide-RNA interaction. As shown in Table 2, evaluations with the RNA-specific scoring function iMDLScore ^*31*^ indicate that ZN423 has the highest rank of binding compared with the four derivatives. This result is consistent with the inability of the derivatives to bind RNA I. Although the score difference itself is not significant for ZN423 and ZN423-4 (e.g., −11.17 vs. −10.48), the loss of activity for ZN423-4 is not surprising because of the possibility of the hydrazide group at R3 to form intra-molecular hydrogen bonds ^*40*^ in addition to the extra size of the hydrazine moiety at R1 position.

**Table 2.** Structure-activity relationship (SAR) of 423 series compounds.

| Name | R<sub>1</sub> | R<sub>2</sub> | R<sub>3</sub> | iMDLScore | Activity |
| --- | --- | --- | --- | --- | --- |
| ZN423 | -H | -H | -CONH<sub>2</sub> | -11.17 | active |
| ZN423-1 | -H | -H | -NHCOCH<sub>3</sub> | -9.99 | inactive |
| ZN423-2 | -H | -H | -CH(CH<sub>3</sub>)CH<sub>3</sub> | -10.32 | inactive |
| ZN423-3 | -H | -CONHNH<sub>2</sub> | -H | -6.64 | inactive |
| ZN423-4 | -NH<sub>2</sub> | -H | -CONHNH<sub>2</sub> | -10.48 | inactive |

## Conclusion

RNAs are historically important therapeutic targets, but they have been less systematically investigated than proteins as targets. X-ray crystallographic and NMR structure studies of ribosomes and rRNA fragments revealed that various aminoglycoside antibiotics inhibit translation by direct binding to architectural elements within the 16S rRNA and demonstrated that RNA can be specifically targeted by small molecules. The array of RNA motifs that can be specifically bound by small molecules and the chemotypes of those small molecules has been expanded through small molecule library-RNA motif library screens^*1, 41*^. However, with few exceptions^*30*^, rational design of chemical inhibitors targeting specific RNA motifs has been less productive due in part to lack of reliable *in silico* and experimental tools for structure-based drug design.

Previously, we evaluated the ability of computational docking to predict experimental binding energies between RNA and small molecules and established a protocol for docking of these complexes^*31*^. However, a significant challenge in accurately docking small molecules to RNA is that the structure of the complexed form of the target RNA is generally unknown. Al-Hashimi and colleagues demonstrated the improved accuracy of *in silico* docking predictions by using multiple conformers of the internal loop region of the TAR RNA hairpin derived from NMR-informed MD simulations ^*30*^. In this study we employed a similar method to screen chemical libraries for small molecule compounds with the potential to specifically bind an RNA containing a UU:GA tandem mismatch. This sequence motif has been predicted to be present in ncRNAs, such as immature miRNAs, implicated in many diseases including cancer. Binding studies with one of the hits identified in our screen, 2-amino-1,3-benzothiazole-6-carboxamide (ZN423), revealed that the motif includes not only the tandem mismatch but also the flanking base pairs. These base pairs contribute to the structural and/or dynamic features necessary to support binding of ZN423. Although not wholly unexpected, the finding that specificity for the UU:GA mismatch is flanking sequence dependent demonstrates the importance of context effect and *increases* the possible number of small non-canonical features that can be specifically targeted. In addition, while the UU:GA tandem mismatch, or more generally YU:GA motif, may appear in multiple RNA molecules, the flanking sequence requirements increase the specificity of ZN423 for only a small subset of RNA molecules.

Finally, ZN423 is small even by small molecule standards and is more akin to a molecular fragment. Thus, the moderate binding affinity (>0.1 mM) for the UU:GA mismatch may be expected to be optimized systematically based on the predicted binding pose within RNA I. The ZN423 series compounds available for testing demonstrates the importance of the carboxamide functionality at position R3, but are less informative of modifications that can be accommodated along the outward-facing opposite edge of ZN423.

## SUPPORTING INFORMATION

Six figures, a table of chemical shifts, and a table listing the top 16 hit compounds are provided.

## ACCESSION NUMBERS

Coordinates have been deposited in the Protein Data Bank under accession number PBD ID: 6VZC. Chemical shifts have been deposited in the Biomolecular Magnetic Resonance Bank under accession number BMRB ID: 50197.

## ACKNOWLEDGEMENTS

We thank Malgorzata Michnicka for preparation of the T7 RNA polymerase and synthesis of the labeled 5’-nucleotide triphosphates.

## REFERENCES

[1] Tran, T., and Disney, M. D. (2011) Molecular recognition of 6’-N-5-hexynoate kanamycin A and RNA 1×1 internal loops containing CA mismatches, Biochemistry 50, 962–969.

[2] Haniff, H. S., Graves, A., and Disney, M. D. (2018) Selective Small Molecule Recognition of RNA Base Pairs, ACS Comb Sci 20, 482–491.

[3] Sztuba-Solinska, J., Shenoy, S. R., Gareiss, P., Krumpe, L. R., Le Grice, S. F., O’Keefe, B. R., and Schneekloth, J. S., Jr. (2014) Identification of biologically active, HIV TAR RNA-binding small molecules using small molecule microarrays, J Am Chem Soc 136, 8402–8410.

[4] Di Giorgio, A., and Duca, M. (2019) Synthetic small-molecule RNA ligands: future prospects as therapeutic agents, Medchemcomm 10, 1242–1255.

[5] Sztuba-Solinska, J., Chavez-Calvillo, G., and Cline, S. E. (2019) Unveiling the druggable RNA targets and small molecule therapeutics, Bioorg Med Chem 27, 2149–2165.

[6] Fulle, S., and Gohlke, H. (2010) Molecular recognition of RNA: challenges for modelling interactions and plasticity, J Mol Recognit 23, 220–231.

[7] Thomas, J. R., and Hergenrother, P. J. (2008) Targeting RNA with small molecules, Chemical reviews 108, 1171–1224.

[8] Disney, M. D., Velagapudi, S. P., Li, Y., Costales, M. G., and Childs-Disney, J. L. (2019) Identifying and validating small molecules interacting with RNA (SMIRNAs), Methods in enzymology 623, 45–66.

[9] Rizvi, N. F., Santa Maria, J. P., Jr., Nahvi, A., Klappenbach, J., Klein, D. J., Curran, P. J., Richards, M. P., Chamberlin, C., Saradjian, P., Burchard, J., Aguilar, R., Lee, J. T., Dandliker, P. J., Smith, G. F., Kutchukian, P., and Nickbarg, E. B. (2019) Targeting RNA with Small Molecules: Identification of Selective, RNA-Binding Small Molecules Occupying Drug-Like Chemical Space, SLAS Discov, 2472555219885373.

[10] Zhang, P., Park, H. J., Zhang, J., Junn, E., Andrews, R. J., Velagapudi, S. P., Abegg, D., Vishnu, K., Costales, M. G., Childs-Disney, J. L., Adibekian, A., Moss, W. N., Mouradian, M. M., and Disney, M. D. (2020) Translation of the intrinsically disordered protein alpha-synuclein is inhibited by a small molecule targeting its structured mRNA, Proc Natl Acad Sci U S A 117, 1457–1467.

[11] Jain, N., Morgan, C. E., Rife, B. D., Salemi, M., and Tolbert, B. S. (2016) Solution Structure of the HIV-1 Intron Splicing Silencer and Its Interactions with the UP1 Domain of Heterogeneous Nuclear Ribonucleoprotein (hnRNP) A1, J Biol Chem 291, 2331–2344.

[12] Bilbille, Y., Vendeix, F. A., Guenther, R., Malkiewicz, A., Ariza, X., Vilarrasa, J., and Agris, P. F. (2009) The structure of the human tRNALys3 anticodon bound to the HIV genome is stabilized by modified nucleosides and adjacent mismatch base pairs, Nucleic Acids Res 37, 3342–3353.

[13] Davlieva, M., Donarski, J., Wang, J., Shamoo, Y., and Nikonowicz, E. P. (2014) Structure analysis of free and bound states of an RNA aptamer against ribosomal protein S8 from Bacillus anthracis, Nucleic Acids Res 42, 10795–10808.

[14] Seok, J. K., Lee, S. H., Kim, M. J., and Lee, Y. M. (2014) MicroRNA-382 induced by HIF-1alpha is an angiogenic miR targeting the tumor suppressor phosphatase and tensin homolog, Nucleic Acids Res 42, 8062–8072.

[15] Wang, H., Yan, B., Zhang, P., Liu, S., Li, Q., Yang, J., Yang, F., and Chen, E. (2020) MiR-496 promotes migration and epithelial-mesenchymal transition by targeting RASSF6 in colorectal cancer, J Cell Physiol 235, 1469–1479.

[16] Yan, T. T., Ren, L. L., Shen, C. Q., Wang, Z. H., Yu, Y. N., Liang, Q., Tang, J. Y., Chen, Y. X., Sun, D. F., Zgodzinski, W., Majewski, M., Radwan, P., Kryczek, I., Zhong, M., Chen, J., Liu, Q., Zou, W., Chen, H. Y., Hong, J., and Fang, J. Y. (2018) miR-508 Defines the Stem-like/Mesenchymal Subtype in Colorectal Cancer, Cancer Res 78, 1751–1765.

[17] Shankar, N., Xia, T., Kennedy, S. D., Krugh, T. R., Mathews, D. H., and Turner, D. H. (2007) NMR reveals the absence of hydrogen bonding in adjacent UU and AG mismatches in an isolated internal loop from ribosomal RNA, Biochemistry 46, 12665–12678.

[18] Davanloo, P., Rosenberg, A. H., Dunn, J. J., and Studier, F. W. (1984) Cloning and Expression of the Gene for Bacteriophage-T7 Rna-Polymerase, P Natl Acad Sci-Biol 81, 2035–2039.

[19] Milligan, J. F., Groebe, D. R., Witherell, G. W., and Uhlenbeck, O. C. (1987) Oligoribonucleotide synthesis using T7 RNA Polymerase and Synthetic DNA Templates, Nucleic Acids Research 15, 8783–8789.

[20] Nikonowicz, E. P., Sirr, A., Legault, P., Jucker, F. M., Baer, L. M., and Pardi, A. (1992) Preparation of C-13 and N-15 Labeled RNAs for Heteronuclear Multidimensional NMR Studies, Nucleic Acids Research 20, 4507–4513.

[21] Denmon, A. P., Wang, J., and Nikonowicz, E. P. (2011) Conformation effects of base modification on the anticodon stem-loop of Bacillus subtilis tRNA(Tyr), J Mol Biol 412, 285–303.

[22] Wang, J., and Nikonowicz, E. P. (2011) Solution structure of the K-turn and Specifier Loop domains from the Bacillus subtilis tyrS T-box leader RNA, J Mol Biol 408, 99–117.

[23] Varani, G., Aboul-ela, F., and Allain, F. H. T. (1996) NMR investigation of RNA structure, Progress in Nuclear Magnetic Resonance Spectroscopy 29, 51–127.

[24] Legault, P., Jucker, F. M., and Pardi, A. (1995) Improved measurement of 13C, 31P J coupling constants in isotopically labeled RNA, FEBS Letters 362, 156–160.

[25] Schwieters, C. D., Kuszewski, J. J., Tjandra, N., and Clore, G. M. (2003) The Xplor-NIH NMR molecular structure determination package, J Magn Reson 160, 65–73.

[26] Yamazaki, T., Muhandiram, R., and Kay, L. E. (1994) Nmr Experiments for the Measurement of Carbon Relaxation Properties in Highly Enriched, Uniformly C-13,N-15-Labeled Proteins - Application to C-13(Alpha) Carbons, J Am Chem Soc 116, 8266–8278.

[27] Lu, X. J., and Olson, W. K. (2003) 3DNA: a software package for the analysis, rebuilding and visualization of three-dimensional nucleic acid structures, Nucleic Acids Res 31, 5108–5121.

[28] Pettersen, E. F., Goddard, T. D., Huang, C. C., Couch, G. S., Greenblatt, D. M., Meng, E. C., and Ferrin, T. E. (2004) UCSF Chimera--a visualization system for exploratory research and analysis, J Comput Chem 25, 1605–1612.

[29] Frank, A. T., Stelzer, A. C., Al-Hashimi, H. M., and Andricioaei, I. (2009) Constructing RNA dynamical ensembles by combining MD and motionally decoupled NMR RDCs: new insights into RNA dynamics and adaptive ligand recognition, Nucleic Acids Res 37, 3670–3679.

[30] Stelzer, A. C., Frank, A. T., Kratz, J. D., Swanson, M. D., Gonzalez-Hernandez, M. J., Lee, J., Andricioaei, I., Markovitz, D. M., and Al-Hashimi, H. M. (2011) Discovery of selective bioactive small molecules by targeting an RNA dynamic ensemble, Nat Chem Biol 7, 553–559.

[31] Chen, L., Calin, G. A., and Zhang, S. (2012) Novel insights of structure-based modeling for RNA-targeted drug discovery, J Chem Inf Model 52, 2741–2753.

[32] Verdonk, M. L., Cole, J. C., Hartshorn, M. J., Murray, C. W., and Taylor, R. D. (2003) Improved protein-ligand docking using GOLD, Proteins 52, 609–623.

[33] Morley, S. D., and Afshar, M. (2004) Validation of an empirical RNA-ligand scoring function for fast flexible docking using Ribodock, J Comput Aided Mol Des 18, 189–208.

[34] Sousa da Silva, A. W., and Vranken, W. F. (2012) ACPYPE - AnteChamber PYthon Parser interfacE, BMC Res Notes 5, 367.

[35] Dieckmann, T., and Feigon, J. (1997) Assignment methodology for larger RNA oligonucleotides: application to an ATP-binding RNA aptamer, Journal of biomolecular NMR 9, 259–272.

[36] Pardi, A. (1995) Multidimensional heteronuclear NMR experiments for structure determination of isotopically labeled RNA, Methods in enzymology 261, 350–380.

[37] Zhang, Q., Sun, X., Watt, E. D., and Al-Hashimi, H. M. (2006) Resolving the motional modes that code for RNA adaptation, Science 311, 653–656.

[38] Clore, G. M., and Schwieters, C. D. (2004) Amplitudes of protein backbone dynamics and correlated motions in a small alpha/beta protein: correspondence of dipolar coupling and heteronuclear relaxation measurements, Biochemistry 43, 10678–10691.

[39] Lee, J., Vogt, C. E., McBrairty, M., and Al-Hashimi, H. M. (2013) Influence of dimethylsulfoxide on RNA structure and ligand binding, Analytical chemistry 85, 9692–9698.

[40] Schonbaum, G. R. (1973) New complexes of peroxidases with hydroxamic acids, hydrazides, and amides, J Biol Chem 248, 502–511.

[41] Childs-Disney, J. L., Tran, T., Vummidi, B. R., Velagapudi, S. P., Haniff, H. S., Matsumoto, Y., Crynen, G., Southern, M. R., Biswas, A., Wang, Z. F., Tellinghuisen, T. L., and Disney, M. D. (2018) A Massively Parallel Selection of Small Molecule-RNA Motif Binding Partners Informs Design of an Antiviral from Sequence, Chem 4, 2384–2404.

