## supporting_information for "2-amino-1,3-benzothiazole-6-carboxamide Preferentially Binds the Tandem Mismatch Motif r(UY:GA)"

### Supporting Tables

#### Table S1. Chemical shifts of RNA I.

| residue | H6/H8 | H2/H5 | H1′ | H2′ | H3′ | H4′ | H5′/5′′ | NH | $NH_2$ |
|---|---|---|---|---|---|---|---|---|---|
| G1 | 8.05 | | 5.74 | 4.81 | 4.42 | | 4.02, 4.17 | | |
| G2 | 7.47 | | 5.84 | 4.57 | 4.52 | 4.46 | 4.43,4.17 | 12.79 | |
| G3 | 7.15 | | 5.69 | 4.46 | 4.36 | 4.44 | 4.41,4.01 | 13.19 | |
| C4 | 7.50 | 5.11 | 5.43 | 4.19 | 4.39 | 4.30 | 4.45,4.01 | | 8.22, 6.82 |
| U5 | 7.74 | 5.36 | 5.46 | 4.62 | 4.48 | 4.33 | 4.46,4.02 | 13.15 | |
| G6 | 7.48 | | 5.65 | 4.43 | 4.18 | 4.42 | 4.36,4.04 | 12.62 | |
| U7 | 7.32 | 5.14 | 5.27 | 4.07 | 4.35 | 4.19 | 4.32,4.97 | 11.06 | |
| G8 | 7.76 | | 5.45 | 4.60 | 4.48 | 4.37 | 4.16,4.06 | | |
| A9 | 8.15 | 7.65 | 5.70 | 4.54 | 4.62 | 4.44 | 4.29,4.17 | | |
| U10 | 7.45 | 5.02 | 5.34 | 4.43 | 4.35 | 4.39 | 4.31,4.17 | 13.19 | |
| G11 | 7.66 | | 5.74 | 4.43 | 4.47 | 4.39 | 4.41,4.08 | 12.60 | |
| C12 | 7.36 | 5.09 | 5.34 | 4.35 | 4.13 | 4.34 | 4.40,3.97 | | 8.46, 6.76 |
| U13 | 7.66 | 5.63 | 5.56 | 3.67 | 4.44 | 4.26 | 4.40,4.00 | 11.66 | |
| U14 | 7.93 | 5.77 | 6.01 | 4.56 | 3.92 | 4.39 | 4.15,3.96 | | |
| C15 | 7.60 | 6.03 | 5.85 | 4.00 | 4.37 | 3.70 | 3.52,2.65 | | |
| G16 | 7.77 | | 5.86 | 4.73 | 5.52 | 4.32 | 4.33,4.10 | 9.86 | |
| G17 | 8.20 | | 4.37 | 4.42 | 4.19 | 4.34 | 4.40,4.16 | 13.28 | |
| C18 | 7.55 | 5.15 | 5.37 | 4.42 | 4.43 | 4.32 | 4.40,3.96 | | 8.53, 6.75 |
| A19 | 7.91 | 7.36 | 5.89 | 4.36 | 4.56 | 4.39 | 4.49,4.06 | | |
| U20 | 7.44 | 5.16 | 5.41 | 4.21 | 4.40 | 4.28 | 4.37,3.99 | | |
| A21 | 7.92 | 7.365 | 5.71 | 4.47 | 4.26 | 4.46 | 4.28,4.05 | | |

| | | | | | | | | | |
|---|---|---|---|---|---|---|---|---|---|
| U22 | 7.37 | 5.12 | 5.07 | 4.13 | 4.22 | 4.29 | 4.28,4.00 | 10.33 | |
| C23 | 7.85 | 5.68 | 5.49 | 4.41 | 4.50 | 4.26 | 4.34,4.03 | | 8.19, 6.90 |
| A24 | 7.98 | 6.99 | 5.83 | 4.61 | 4.58 | 4.47 | 4.45,4.12 | | |
| G25 | 7.27 | | 5.59 | 4.32 | 4.40 | 4.38 | 4.42,4.01 | 13.37 | |
| C26 | 7.55 | 5.08 | 5.40 | 4.26 | 4.42 | 4.31 | 4.46,3.98 | | 8.42, 6.76 |
| C27 | 7.70 | 5.40 | 5.43 | 4.34 | n.a. | 4.34 | 4.37,3.97 | | 8.45, 6.76 |
| C28 | 7.61 | 5.43 | 5.67 | 3.92 | 4.12 | 4.09 | 4.40,3.94 | | 8.17, 7.01 |

The non-exchangeable $^{1}$H chemical shifts were measured at 26 °C and pH 6.8 and are referenced with DSS set to 0.00 ppm. Exchangeable proton chemical shifts were measured at 15 °C. The uncertainties in the chemical shift values are ≈0.015 ppm. The 5' and 5'' protons are not stereospecifically assigned.

**Table S1. (continued)**

| residue | C6/C8 | C2/C5 | C1’ | C2’ | C3’ | C4’ | C5’ | N(H) | A/C N6/N4 |
|---|---|---|---|---|---|---|---|---|---|
| G1 | 139.20 | | | - | - | - | - | | - |
| G2 | 136.94 | | 91.90 | 74.85 | 72.20 | 81.44 | 65.18 | 147.84 | |
| G3 | 136.11 | | 92.22 | 74.45 | 72.27 | 81.70 | 64.85 | 148.95 | |
| C4 | 141.19 | 96.37 | 93.45 | 74.68 | 71.28 | 81.05 | 63.69 | | 97.52 |
| U5 | 141.83 | 103.10 | 92.74 | 74.78 | 71.81 | 81.11 | 63.94 | 161.67 | |
| G6 | 136.12 | | 92.09 | 74.58 | 72.27 | 81.31 | 65.50 | 147.50 | |
| U7 | 140.3`3 | 102.97 | 92.89 | 74.71 | 71.41 | 82.03 | 63.52 | 158.30 | |
| G8 | 139.42 | | 88.40 | 74.78 | 71.81 | 81.18 | 66.18 | | |
| A9 | 141.55 | 154.11 | 89.83 | 74.45 | 73.99 | 82.83 | 66.17 | | |
| U10 | 141.96 | 102.84 | 92.35 | 74.19 | 72.80 | 81.97 | 65.67 | 162.11 | |
| G11 | 136.66 | | 91.71 | 74.58 | 72.00 | 81.18 | 64.77 | 147.79 | |
| C12 | 140.69 | 96.41 | 93.00 | 74.52 | 71.02 | 81.05 | 63.77 | | 99.25 |
| U13 | 140.88 | 104.27 | 93.65 | 75.04 | 72.27 | 81.64 | 63.11 | 159.94 | |
| U14 | 144.91 | 104.65 | 88.34 | 73.66 | 76.89 | 86.13 | 66.92 | | |
| C15 | 143.05 | 97.66 | 88.15 | 76.76 | 79.46 | 83.62 | 66.42 | | |
| G16 | 143.14 | | 93.71 | 76.30 | 75.41 | 82.37 | 68.10 | 143.86 | |
| G17 | 139.06 | | 92.35 | 74.89 | 74.37 | 83.62 | 68.22 | 148.32 | |
| C18 | 141.19 | 96.37 | 93.07 | 74.32 | 71.15 | 80.98 | 63.35 | | 97.45 |
| A19 | 139.83 | 153.65 | 92.16 | 74.98 | 71.94 | 81.25 | 64.06 | | |
| U20 | 141.46 | 102.97 | 91.64 | 74.91 | 72.60 | 81.05 | 64.35 | | |
| A21 | 140.79 | 154.06 | 91.51 | 74.58 | 73.66 | 83.02 | 66.18 | | |
| U22 | 142.33 | 102.57 | 92.68 | 74.00 | 71.94 | 81.01 | 64.27 | 156.71 | |
| C23 | 142.19 | 97.47 | 92.03 | 74.71 | 72.33 | 81.97 | 64.22 | | 97.81 |
| A24 | 140.06 | 152.93 | 92.16 | 74.71 | 72.80 | 81.57 | 65.43 | | |
| G25 | 136.03 | | 91.77 | 74.45 | 71.81 | 81.11 | 64.52 | 148.51 | |
| C26 | 141.19 | 96.24 | 93.13 | 74.65 | 72.47 | 81.71 | 64.72 | | 98.49 |
| C27 | 141.65 | 96.95 | 93.22 | 74.71 | | 81.01 | 63.61 | | 98.29 |
| C28 | 142.10 | 97.28 | 91.90 | 76.70 | 68.84 | 82.63 | 64.35 | | 96.92 |

The $^{13}$C frequency corresponding to 0.00 ppm was set according to the 0.00 ppm $^{1}$H (DSS) frequency at 26 °C using the ratio $\gamma_C/\gamma_H$. The chemical shifts have uncertainties of ≈0.075 ppm. The $^{15}$N frequency corresponding to 0.00 ppm was set according to the 0.00 ppm $^{1}$H (DSS) frequency at 15 °C using the ratio $\gamma_N/\gamma_H$. The chemical shifts have uncertainties of ≈0.055 ppm.

**Table S1. (continued)**

| residue | YN1/ RN9 | YC2/ GC2 | YC4/ GC6 | RN7 | A N1 | A N3 |
|---|---|---|---|---|---|---|
| G1 | - | - | - | - | - | - |
| G2 | 170.14 | 156.67 | 161.79 | 234.48 | | |
| G3 | 169.54 | 157.12 | 161.90 | 234.78 | | |
| C4 | | 159.29 | | | | |
| U5 | | 152.92 | 169.70 | | | |
| G6 | 170.51 | 156.21 | 161.67 | 237.00 | | |
| U7 | | 154.27 | 166.87 | | | |
| G8 | 172.44 | 160.2 | | 231.41 | | |
| A9 | 169.47 | | | 231.37 | 223.94 | 215.04 |
| U10 | | 152.87 | | | | |
| G11 | 170.58 | 156.69 | 161.54 | 235.30 | | |
| C12 | | 158.90 | | | | |
| U13 | | 153.31 | | | | |
| U14 | | 155.30 | | | | |
| C15 | | 160.30 | | | | |
| G16 | 167.46 | 155.86 | 161.77 | 237.68 | | |
| G17 | 171.1 | 157.22 | 162.60 | 233.81 | | |
| C18 | | 159.11 | | | | |
| A19 | 171.55 | | | 231.81 | 222.83 | 214.37 |
| U20 | | 153.36 | | | | |
| A21 | 169.69 | | | 231.89 | 226.24 | 216.30 |
| U22 | | 152.25 | 168.23 | | | |
| C23 | | 159.35 | | | | |
| A24 | 170.80 | | | 231.22 | 220.75 | 213.85 |
| G25 | 169.76 | 157.27 | 161.67 | 234.63 | | |
| C26 | | 159.11 | | | | |
| C27 | | 159.29 | | | | |
| C28 | | 160.22 | | | | |

The $^{13}C$ frequency corresponding to 0.00 ppm was set according to the 0.00 ppm $^{1}H$ (DSS) frequency at 26 °C and 15 °C using the ratio $\gamma_C/\gamma_H$. The chemical shifts have uncertainties of ≈0.075 ppm. The $^{15}N$ frequency corresponding to 0.00 ppm was set according to the 0.00 ppm $^{1}H$ (DSS) frequency at 26 °C using the ratio $\gamma_N/\gamma_H$. The chemical shifts have uncertainties of ≈0.055 ppm.

### Table S2: The selected 16 hits from virtual screening.

| NO. | Mol Name | Structure | Catalog ID | Mol Weight |
|---|---|---|---|---|
| 1 | 2-[(4-chlorophenyl)(1,1-dioxido-1,2-benzisothiazol-3-yl)amino]ethanol | 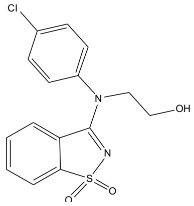 | ZN867 | 336.8 |
| 2 | 7-(2-fluorobenzyl)-3-methyl-8-piperazin-1-yl-3,7-dihydro-1H-purine-2,6-dione | 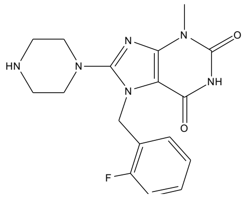 | ZN119 | 358.4 |
| 3 | 1-deoxy-1-[(3,4-dimethylphenyl)amino]pentitol | 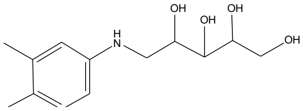 | ZN890 | 255.3 |
| 4 | 4-(2-hydroxyphenyl)-6-methyl-N-(2-methylphenyl)-2-thioxo-1,2,3,4-tetrahydro-5-pyrimidinecarboxamide | 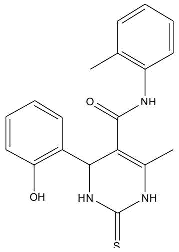 | ZN591 | 353.4 |
| 5 | 3-[(pyridin-3-ylmethyl)amino]-1,2,4-triazin-5(4H)-one | 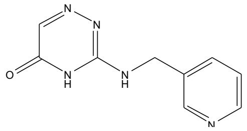 | ZN179 | 203.2 |
| **6** | **2-amino-1,3-benzothiazole-6-carboxamide** | 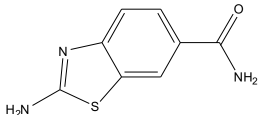 | **ZN423** | **193.2** |
| 7 | methyl 4-{[2-(acetylimino)-4-oxo-1,3-thiazolidin-5-ylidene]methyl}benzoate | 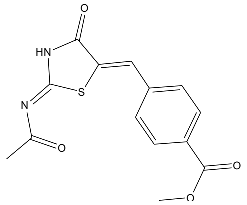 | ZN759 | 304.3 |
| 8 | 2-chloro-5-(1-piperidinylsulfonyl)-N-(2-thienylmethyl)benzamide | 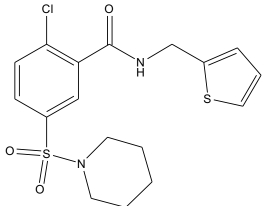 | ZN719 | 398.9 |
| 9 | 2-{[(6-bromo-1,3-benzodioxol-5-yl)methylene]amino}-4,6-dimethylphenol | 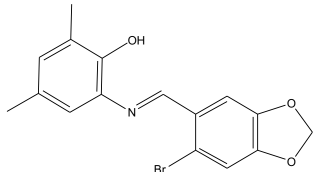 | ZN812 | 348.2 |

| | | | | |
|---|---|---|---|---|
| 10 | 4-(1H-benzimidazol-2-yl)-1,2,5-oxadiazol-3-amine | 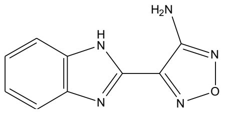 | ZN435 | 201.2 |
| 11 | 2-{[(4-amino-2-methyl-5-pyrimidinyl)methyl]thio}-6-methyl-4(3H)-pyrimidinone | 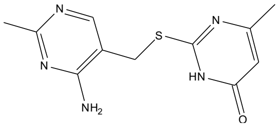 | ZN681 | 263.3 |
| 12 | 2-[(4-methylbenzyl)thio]-4(3H)-pyrimidinone | 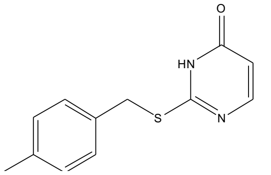 | ZN620 | 232.3 |
| 13 | N~2~-(2-furylmethyl)-N~1~-1-naphthyl-alpha-asparagine | 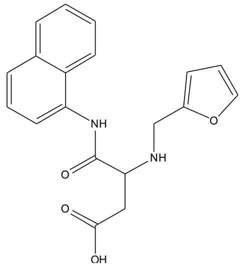 | ZN183 | 338.4 |
| 14 | 3-{[2-(4-morpholinyl)-2-oxoethyl]thio}-1H-1,2,4-triazol-5-amine | 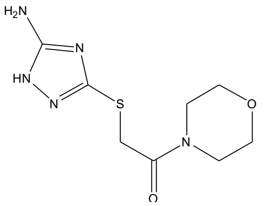 | ZN957 | 243.3 |
| 15 | 8-methyl-3-(methylthio)-5H-[1,2,4]triazino[5,6-b]indole | 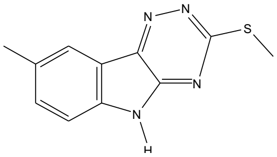 | ZN629 | 230.3 |
| **16** | **5,7-dimethyl-1,4-dihydro-2,3-quinoxalinedione** | 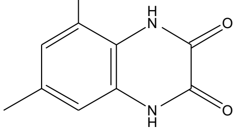 | **ZN449** | **190.2** |

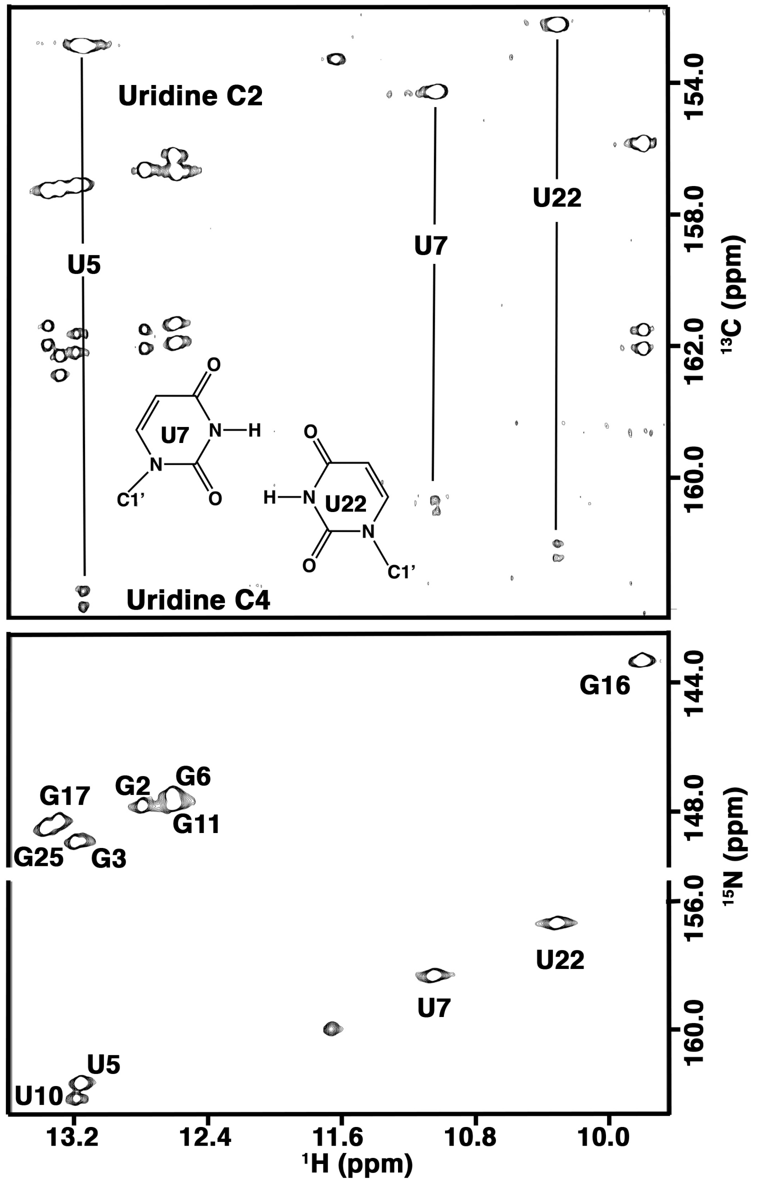


**Figure S1.** Imino proton regions of H(N)CO (top) and $^{15}N$-$^{1}H$ HSQC (bottom) spectra. The U20 H3 is broadened by solvent exchange and does not appear in the spectra. The downfield shifted C2 and C4 resonances of U7 and U22, respectively, are indicative of participation of the attached oxygen atoms in a hydrogen bond.

**A.**

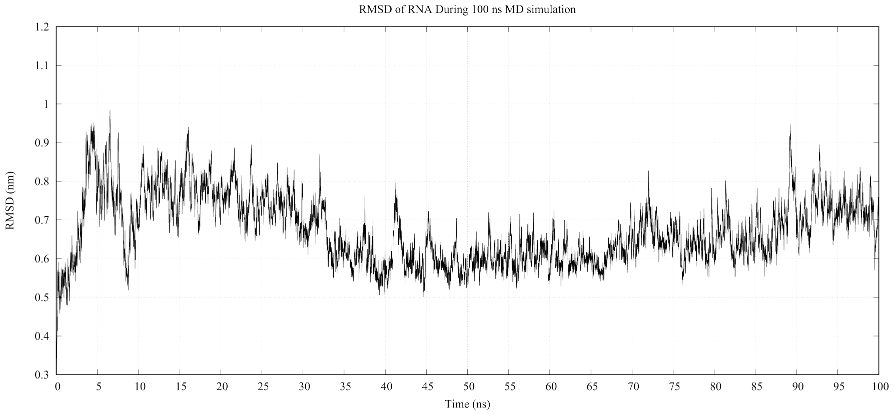


**B.**

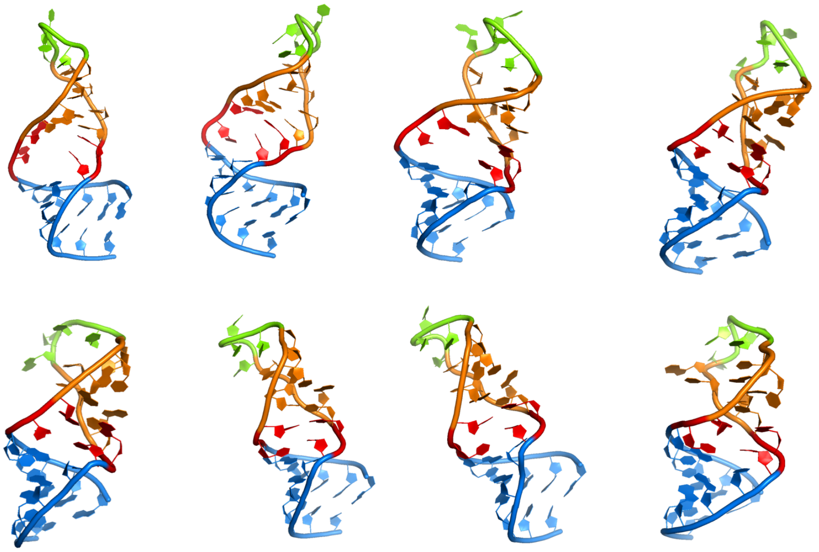

**Figure S2.** Molecular dynamics simulation and the structural ensemble selection. A. The trajectory of MD simulation. B. 8 of 20 selected RNA I structures based on experimental NMR RDC data as the representative ensemble.

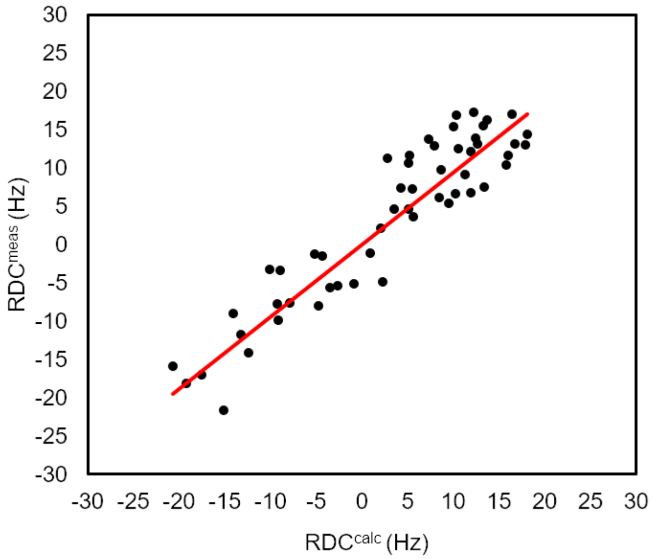


**Figure S3:** Comparison of in silico model and experimental RNA I structure. Corrrelation of experimental RDC and calculated RDC data. $R^2$=0.86 and RMSD=4.12Hz.

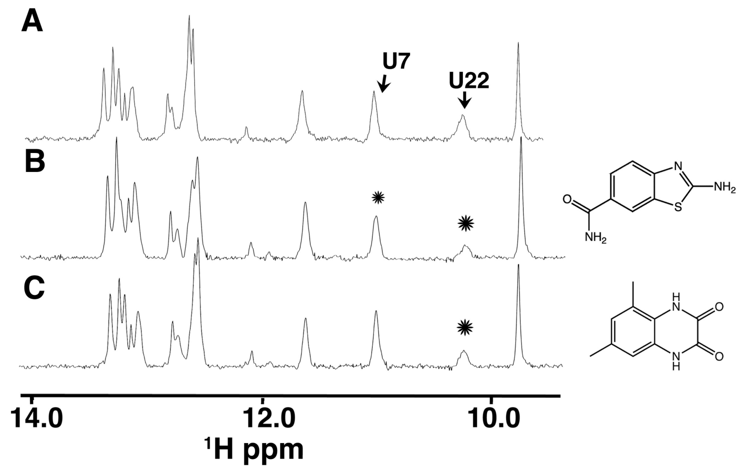


**Figure S4.** 1D $^1$H NH spectra of the GA:UU mismatch RNA hairpin. (A) RNA (0.015 mM) in 5% DMSO, (B) with ZN423, and (C) 5,7-dimethyl-1,4-dihydro-2,3-quinoxalinedione. Peaks altered by the compounds are labeled (*). Of the two compounds, ZN423 was found to interact more strongly with RNA I and so was selected for additional. Compound concentrations are 0.1 mM.

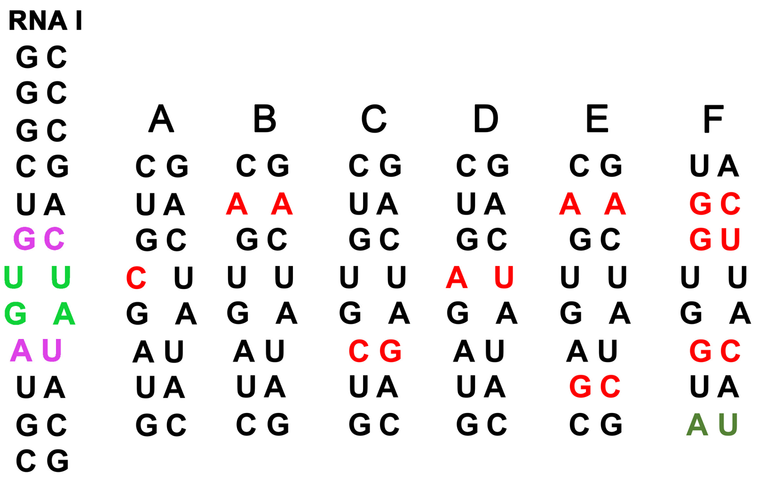


**Figure S5.** Nucleotide substitutions in RNA I and the effect on ZN423 binding as assessed by NMR spectral perturbation. (A) Displays spectral properties similar to RNA I indicative of similar binding affinity. The spectral perturbations displayed by (B) in the presence ZN423 are indicative of a weak interaction. Molecules containing the substitutions C-F display no N-H or base C-H NMR spectral evidence of interaction.

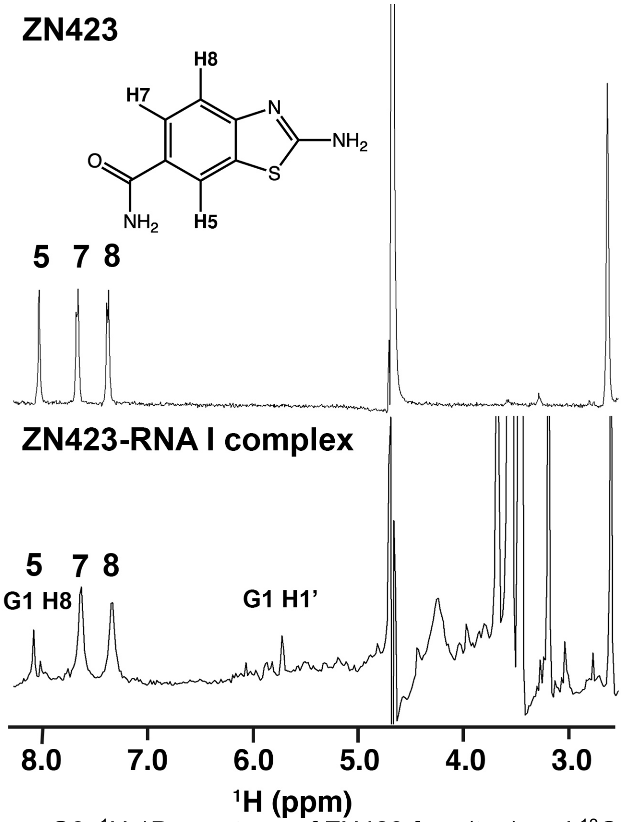


**Figure S6**. $^{1}$H 1D spectrum of ZN423 free (top) and $^{13}$C-filtered $^{1}$H 1D spectrum of ZN423 in the presence of RNA I. The G1 of RNA I was ca. 25% $^{13}$C labeled and so protons corresponding to G1 are incompletely filtered from the lower spectrum. Even though the G1 H8 and ZN423 H5 resonances are nearly overlapped, the ZN423 H5 is clearly exchange broadened, consistent with its positioning deep in the minor groove. The same exchange phenomenon that causes broadening of the RNA resonances also causes broadening of the ZN423 H5.
